# A low-cost, long-term underwater camera trap network coupled with deep residual learning image analysis

**DOI:** 10.1101/2021.03.08.434472

**Authors:** Stephanie M. Bilodeau, Austin W. H. Schwartz, Binfeng Xu, V. Paúl Pauca, Miles R. Silman

**Affiliations:** Department of Biology, Wake Forest University, 1843 Wake Forest Rd, Winston-Salem, NC 27109; Center for Energy, Environment, and Sustainability, Wake Forest University, 1843 Wake Forest Rd, Winston-Salem, NC 27109; Department of Computer Science, Wake Forest University, 1843 Wake Forest Rd, Winston-Salem, NC 27109

**Keywords:** Behavior, camera trap, image classification, long-term, machine learning, marine, underwater, reefscape, deep learning, landscape of fear

## Abstract

1. Understanding long-term trends in marine ecosystems requires accurate and repeatable counts of fishes and other aquatic organisms on spatial and temporal scales that are difficult or impossible to achieve with diver-based surveys. Long-term, spatially distributed cameras, like those used in terrestrial camera trapping, have not been successfully applied in marine systems due to limitations of the aquatic environment.
2. Here, we develop methodology for a system of low-cost, long-term camera traps (**D**ispersed **E**nvironment **A**quatic **C**ameras), deployable over large spatial scales in remote marine environments. We use machine learning to classify the large volume of images collected by the cameras. We present a case study of these combined techniques’ use by addressing fish movement and feeding behavior related to grazing halos, a well-documented benthic pattern in shallow tropical reefscapes.
3. Cameras proved able to function continuously underwater at deployed depths (up to 7 m, with later versions deployed to 40 m) with no maintenance or monitoring for over five months, and collected time-lapse images during daylight hours for a total of over 100,000 images. Our ResNet-50-based deep learning model achieved 92.5% overall accuracy in sorting images with and without fish, and diver surveys revealed that the camera images accurately represented local fish communities.
4. The cameras and machine learning classification represent the first successful method for broad-scale underwater camera trap deployment, and our case study demonstrates the cameras’ potential for addressing questions of marine animal behavior, distributions, and large-scale spatial patterns.

## Introduction

Terrestrial camera trapping is a growing field and a technique increasingly applied in global biodiversity monitoring (e.g., Steenweg et al. 2017). Camera traps are remotely activated cameras that rely on motion and heat sensors to trigger when an animal passes by. They are used to study species richness (Rowcliffe 2017) and the distribution, abundance, habitat use, and behavior of wildlife around the world, with many studies surveying more than one species at a time (Burton et al. 2015). Most camera traps are small, relatively inexpensive, and often deployed in groups or networks over a wide area for months at a time. They are typically less invasive and more reliable than comparable observation techniques (Cutler and Swann 1999). Because they trigger based on the difference between background radiation and a warm-bodied animal passing through the sensor’s field, camera traps have typically been used to study mammals and birds, although the field is now expanding to include some ectotherms (Rowcliffe 2017).

Comparable techniques for monitoring underwater species face several technical challenges, chiefly the attenuation of infrared radiation in water, which renders a standard commercial camera trap unlikely to trigger underwater, except at very close range (Giles and Bankman 2005), and the rigors of operating in the marine environment where water intrusion and algal and faunal fouling are persistent issues. The fact that most fish are ectotherms further complicates use of the traditional heat-triggered infrared sensor technology used in most terrestrial camera traps. Far-red illumination invisible to most fish provides one potential alternative to infrared (Williams et al. 2014), although far-red light still attenuates over short distances underwater. Given the ability of sound to propagate well underwater, acoustic techniques provide another possible alternative to infrared sensing (Giles and Bankman 2005). Acoustic cameras have been used in the past to image sharks and other fish in low-light, turbid environments, replacing light-based imaging entirely (McCauley et al. 2016). This suggests that acoustic techniques could also be used to trigger conventional optical cameras.

Due to the power requirements of active triggering (far-red light, sonar) and recording methods, current underwater fish monitoring and measurement techniques are either limited by short battery life and operate on the scale of hours, as with baited remote underwater video (BRUV) and similar short-term recording devices (Cappo et al. 2004, Colton and Swearer 2010, Brooks et al. 2011, Williams et al. 2014, Boussarie et al. 2016) or require a tethered, external power source (Boom et al. 2014, Marini et al. 2018). Because of this, the spatial extent, number of cameras, and duration of monitoring for marine systems are vastly smaller in scope than terrestrial efforts (compare Williams et al. 2014 and Siddiqui et al. 2018 to the TEAM or Snapshot Serengeti datasets described in Beaudrot et al. 2016, Norouzzadeh et al. 2018, respectively). Even long-term underwater monitoring with an external power supply may be limited to recording images or video during daylight hours (e.g., Boom et al. 2014) due to the difficulties of avoiding reflected particulate matter in underwater images taken at night with direct illumination. Without a side-mounted or similarly external flash, both white light and infrared images will be obscured by the illumination of biotic and abiotic particulates suspended in the water column, and continuous illumination may also attract fish and other marine organisms, depending on the color of the light (Ko et al. 2018). Thus, most long-term underwater observations are limited with regard to both power and nighttime illumination, and solutions can be costly and prohibit deployment of underwater cameras in remote locations without access to external power or consistent upkeep.

A related challenge faced by both terrestrial and marine camera traps is the time cost related to processing and analyzing large photosets obtained from multiple cameras over the course of months or years (Norouzzadeh et al. 2018). While computer vision techniques are not yet widespread in the field of camera trapping, they have the potential to reduce time cost (Rowcliffe 2017) and have already been successfully applied to the extensive Snapshot Serengeti camera trap dataset (Norouzzadeh et al. 2018) as well as video frames from cabled or short-term marine cameras (Boom et al. 2014, Siddiqui et al. 2018, Marini et al. 2018, Villon et al. 2018).

Here we present a simple design for an affordable, long-running, autonomous underwater camera based on an existing commercially-available terrestrial model. We outline the deployment of our Dispersed Environment Aquatic Cameras (DEACs) across a 270 km^2^ tropical reefscape, our testing, and our subsequent analysis of the 100,000+ images obtained by implementing a deep convolutional neural network (CNN) technique for image classification. To demonstrate both the efficacy of our design and the value of long-term unmanned underwater observations to marine ecology research, we present a case-study in fish feeding behavior as it relates to a well-documented benthic pattern at Lighthouse Reef Atoll, Belize.

## Methods

### Design

#### Camera Selection

Cuddeback Silver Series scouting cameras, model 1231 (Cuddeback, Green Bay, WI, USA), were chosen for their compact size, their relatively high 20 megapixel (MP) image resolution, and their time-lapse function, which allows images to be taken at pre-programmed time intervals and certain light levels without requiring the use of additional video or motion-triggered image settings. This programming flexibility, especially regarding the time-lapse feature, is not included in multiple similar cameras from other manufacturers. The cameras were programmed to take 20 MP images every 15 minutes whenever ambient light levels were high enough to allow for color photography without flash, using the “Day” setting. Nighttime images and video and all infrared sensor-triggered images and video were disabled to conserve both power and memory space because initial field tests showed that few or no additional usable images were captured using the infrared, low-light “Night” setting.

Cameras were synchronized to record photos on the hour and every 15 minutes following to ensure that images from different cameras and sites were captured at the same time of day and under the same local conditions. The 15-minute interval was chosen to balance the need for regular observations of reef and seagrass communities that may include transient fish species and the constraints of storing and processing thousands of high-resolution images collected by multiple cameras over a months-long deployment. The ability of this interval to adequately capture the community composition and species present at a given reef was validated using in-person diver surveys (described below).

Memory cards were 32 GB SanDisk (Western Digital Corporation, Milpitas, CA, USA) or Kingston (Kingston Technology Corporation, Fountain Valley, CA, USA) microSD cards with adapters, capable of holding over 25,000 20 MP color images each. Each camera required eight Energizer Ultimate Lithium AA batteries (Energizer Holdings, Inc., St. Louis, MO, USA), which provided enough power for over five months of continuous function under the settings described here.

The total cost of each camera, including the batteries and SD card, came to just $125 per unit with the housing (discussed below). The use of pre-built commercial trail cameras, which are designed for energy efficiency over long deployments, significantly reduced both material and energy costs, relative to constructing a similar camera from scratch using components like a Raspberry Pi (Raspberry Pi Foundation, Cambridgeshire, UK) and GoPro (GoPro, Inc., San Mateo, CA, USA), Canon (Canon, Inc., Ota City, Tokyo, Japan), or Sony (Sony Corporation, Minato City, Tokyo, Japan) cameras, as used in previous underwater camera applications (e.g., Brooks et al. 2011, Williams et al. 2014, Boussarie et al. 2016, Siddiqui et al. 2018, Villon et al. 2018).

#### Housing Construction

Two different housings were tested in the field, both based on commercially available junction box enclosures. Housing 1, based on item DS-AT-1217-1 available from “Saipwell” (Saip Electric Group Co., Ltd, Wenzhou, China), has thinner walls (2.4 - 3.9 mm) made of an unspecified plastic. The top secures with specially-shaped plastic screws and it contains an O-ring like insert made of foam. Housing 2, based on model ML-47F^*^1508 from Polycase, Inc. (Avon, OH, USA), is a thick-walled (3.5-4.0 mm) design made of polycarbonate resin secured with stainless steel screws and a silicon rubber gasket. Both housings had a 2 inch diameter hole drilled in the faceplate and a 3 inch disk of 1/8 inch thick acrylic epoxied to the opening with MarineWeld (J-B Weld Company, Atlanta, GA, USA) to act as a window. Since both housings used identical acrylic windows and contained the same cameras, images collected using each design were indistinguishable; the chief difference between the housings was their pressure tolerance and leakage at depth. Each housing cost approximately $30 for all components (included in the total price given above). Ablative antifouling boat paint was applied to the housing exterior, excluding the back and acrylic window, of a subset of cameras with Housing 2 to reduce biofouling.

Cameras were programmed, armed, and packed inside their housings with cardboard spacers. Housings were sealed with Star brite marine silicone sealant (Star brite, Fort Lauderdale, FL, USA) around the seam of the enclosure lid. Silicone sealant was also used to reinforce the edges of the epoxy seal around the lens window, both inside and outside.

#### Field Installation

Cameras were secured to four-legged bent rebar stands with plastic cable ties threaded through mounting holes pre-built into the housings (Fig 1). Due to their buoyancy, camera housings were placed underneath the crossed rebar forming the top of each stand and secured laterally to the four legs, which were sunk into the sediment to keep the stands upright. Lightweight plastic or polystyrene buoys were tethered to a small subset of cameras and stands located in particularly shallow water (2 m or less) to prevent collision by boats.

**Figure 1.**
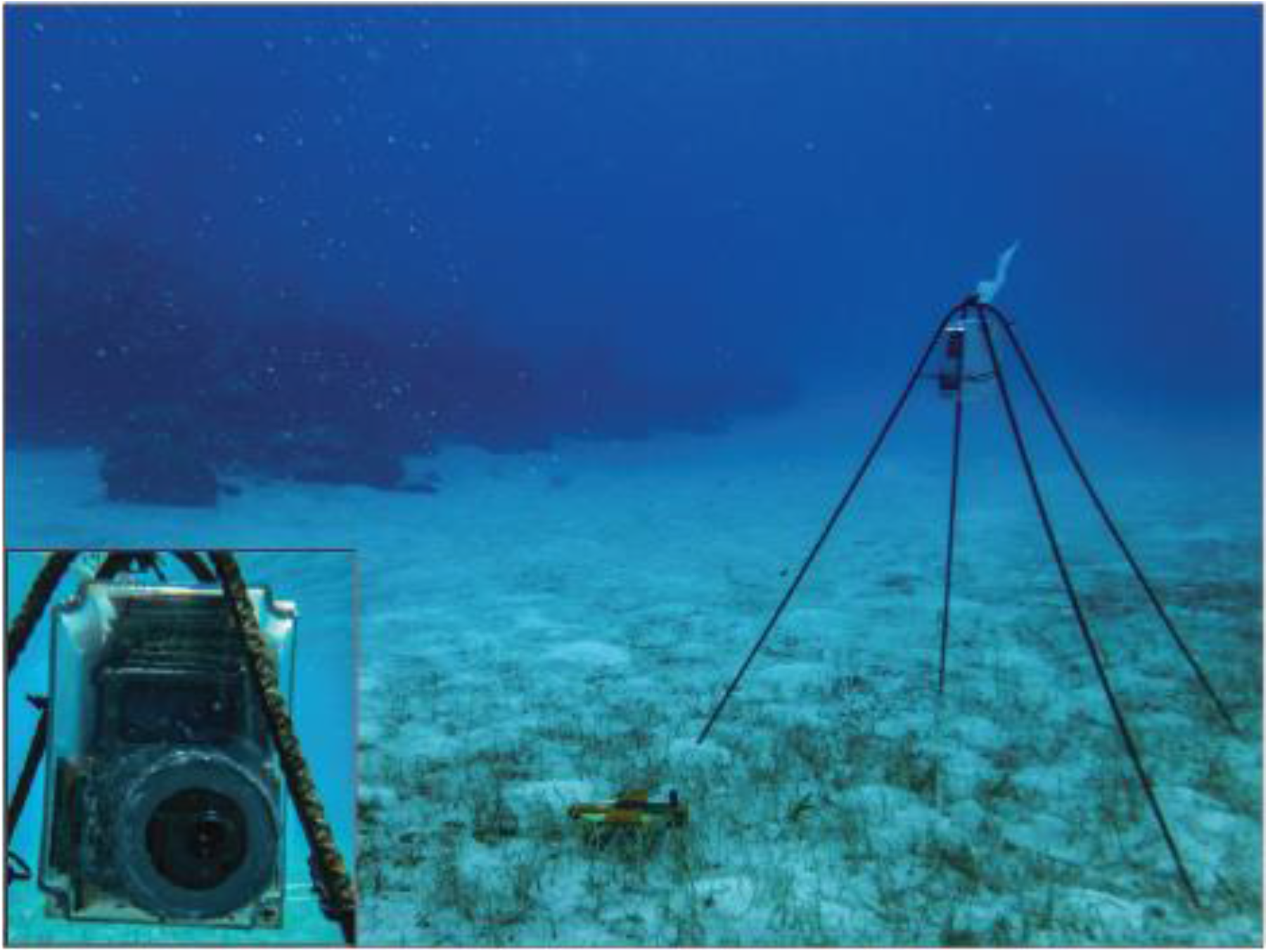
DEACs were deployed on four-legged rebar stands in the field. Plastic cable ties attached to the camera housing and to the legs were used to secure buoyant cameras in housings below the crossed rebar at the top of each stand. Bottom left inset: A detailed front view of a camera in Housing 1 underwater.

### Deployment

#### Study System and Question

Lighthouse Reef Atoll off the coast of Belize is primarily a shallow (1-8 m depth) lagoon environment, dominated by scattered patch reefs interspersed with a mosaic of seagrass, macroalgae, and sand. Although relatively isolated from the mainland, the atoll is subject to heavy local fishing pressure for conch, lobster, and certain fish species. A small no-take marine protected area (MPA) surrounds the island of Half Moon Caye in the southeastern corner of the atoll. Because of its shallow benthos, wide variety of benthic cover types, and spatial variation in protected status, Lighthouse Reef provides an ideal location to test our DEACs under a variety of conditions. The shallow marine grazing system of Lighthouse Reef and similar Caribbean locations is in many ways analogous to terrestrial grazing systems like the African savanna (Burkepile 2013), where camera trap networks have been effectively deployed for years (e.g., Norouzzadeh et al. 2018).

To structure our testing and demonstrate the utility of the DEACs for addressing ecological questions at large spatial scales, we organized our deployments around a widespread benthic pattern particularly prominent at Lighthouse Reef: grazing halos. Grazing halos consist of a bare sand or lightly vegetated border surrounding a coral patch or similar underwater structure that separates the reef from surrounding dense vegetation (i.e. seagrass or algae). The heightened grazing observed inside these halos could be due to a landscape of fear (e.g., Hammerschlag et al. 2015), where herbivores are afraid to venture past a threshold distance from the reef due to predation risk (Madin et al. 2011). At Lighthouse Reef, grazers are mostly large parrotfishes (Scaridae) and surgeonfishes (Acanthuridae). However, this threshold of fish density could also be due simply to the natural dispersion of grazers as they venture farther from the reef, which serves as a central aggregating structure for many fish (Sale and Douglas 1984, Bohnsack 1989, Layman et al. 2013). We proposed that if a strong landscape of fear is in effect, herbivorous species observed and photographed in the halo will never venture out into the surrounding seagrass, except perhaps when traveling in schools. However, a simple drop-off in fish density with distance would still result in the occasional grazing reef fish being seen by our cameras, and if reef fishes are food limited rather than predator limited, they should forage widely in the seagrass, which is a preferred food (Bilodeau 2019). Therefore, regular detection of reef herbivores out in the seagrass over multiple months could be taken as evidence refuting the landscape of fear at Lighthouse Reef.

#### Spatial Arrangement

Cameras were deployed in pairs at 21 patch reef sites within Lighthouse Reef Atoll. DEAC sites were distributed evenly inside and outside of the Half Moon Caye Natural Monument MPA in the southeastern corner of the atoll. Seven sites (3 inside the MPA, 4 outside) featured predominantly algal bottom cover; the rest were in areas surrounded by seagrass, primarily *Thalassia testudinum*. Patch reef sites for camera deployment were chosen via random point placement using satellite imagery and a depth map of the atoll (courtesy of the Carnegie Airborne Observatory), which allowed sites to be evenly stratified across depths from 2-7 m.

Sites deeper than 4 m were initially avoided due to leakage of Housing 1 past this depth, although depths of up to 7 m (maximum depth required in the study) were successfully achieved with Housing 2, which was used for initial deployments in March 2018 and all deployments from August 2018 onward. Cameras were deployed in sets of two, one camera located at the edge of the sandy halo surrounding a patch reef and the other located at least twice the halo’s width away from the edge in the surrounding seagrass or algal benthic cover (Fig 2). In the case of particularly narrow halos, “control” cameras were placed a minimum of 15 m from the halo edge. This allowed control cameras a view of the same bottom cover (seagrass or macroalgae) as that adjacent to the halo but placed them well beyond the fish density thresholds observed by Layman et al. (2013), while also accounting for the possibility that larger halos could represent reefs with larger or farther-ranging fish populations. “Halo” cameras had a relatively wide field of view, as is typical of terrestrial trail cameras, which included both the halo in front of them and the patch reef beyond. Both cameras were pointed toward the reef, although the edge of the halo was beyond the range of view for the grass (“control”) camera at most sites.

**Figure 2.**
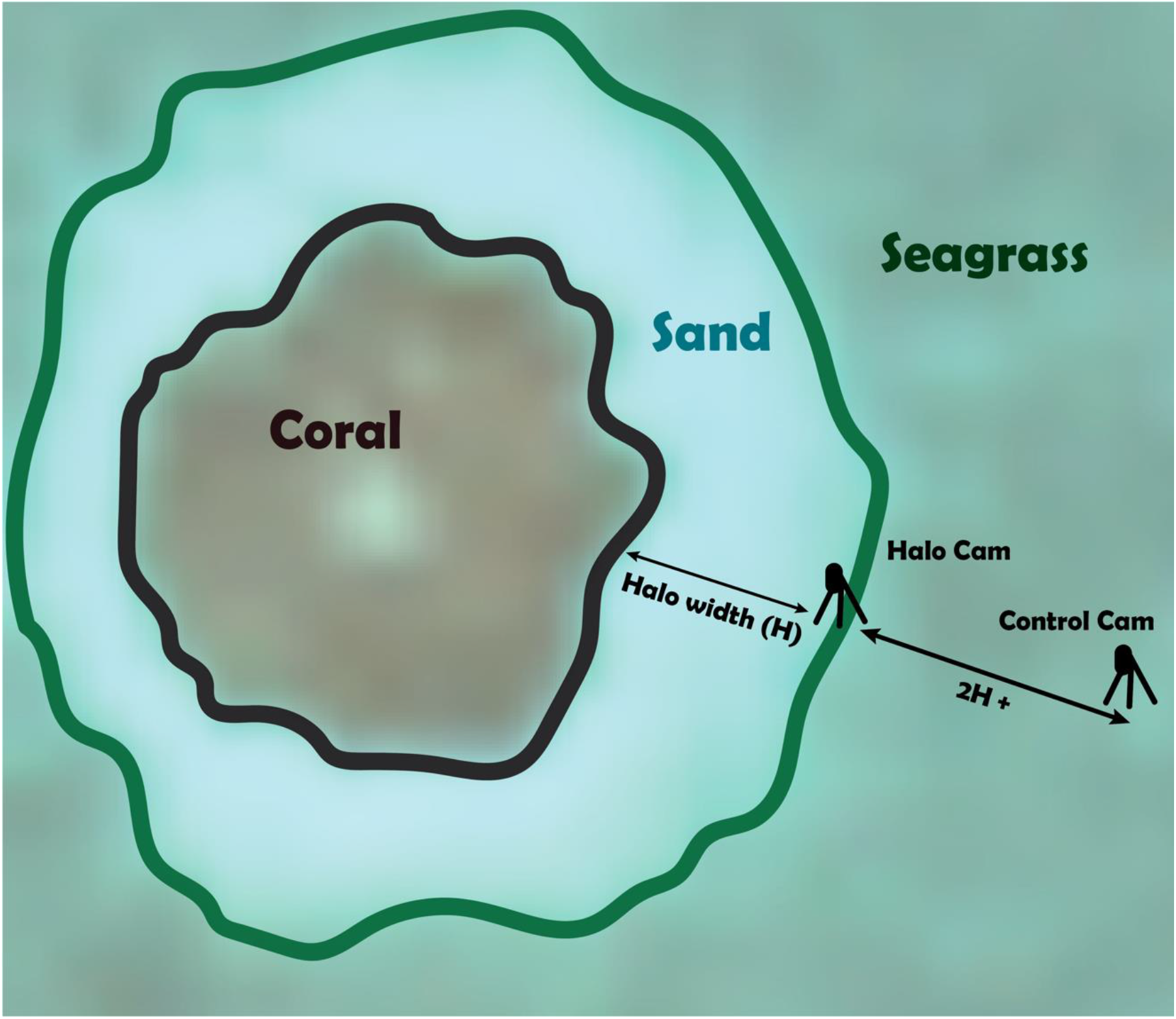
Camera arrangement at a patch reef site. One camera was placed on the edge of the halo facing in toward the reef (“Halo Cam”), while the second (“Control Cam”) was placed twice the halo’s width away in the surrounding seagrass or algal cover, in line with the first camera.

#### Monitoring and Secondary Deployments

Camera deployments occurred in stages, with the first cameras deployed in March 2018 and the last cameras collected in March 2019 (Fig S1). The longest-running DEACs remained continuously active underwater from August 2018 to January 2019, a five-month interval, and were still operating at retrieval. Initial deployment of cameras in March 2018 included one camera at the edge of a halo and a second camera pointed at the first in order to assess whether the presence of the camera and stand had any effect on the presence or behavior of marine animals. Given that no effects of camera presence were noted in this initial test, doubling of cameras in this manner was discontinued for future deployments, with the two cameras at each site positioned to monitor different benthic environments and unable to see each other. During the summer of 2018 (June to August), cameras were consistently checked every 1-2 weeks for leakage, algal overgrowth, or stand displacement. Cameras deployed at longer intervals from March to June 2018, August 2018 to January 2019, and January to March 2019 were unmonitored during these periods in order to test long-term underwater function and determine the effects of biofouling on housings and image quality in the absence of cleaning or regular adjustments.

#### Diver Observations

In-water observations were conducted by a team of 2-3 divers at each camera site and at several additional locations in order to validate the camera’s ability to detect fish and other animals. Observations consisted of all divers sitting directly behind each camera for 15 minutes and recording the presence and abundance of all fish and other animals observed on the reef, in the halo, or in the grass to the genus or species level, when possible. Divers faced forward toward the reef and recorded all fish observed within the halo or on the reef itself, as this was the primary field of view of the camera. In the seagrass or macroalgae, divers again faced toward the reef and recorded only fish that swam in front of them and the camera. Divers remained stationary for the entire observation period at each camera, and their presence did not have any observable effects on fish within the camera’s view, with multiple fish of different species swimming quite close to divers during the observations. Fish species and counts were determined by a consensus of all the divers present at each observation. A non-metric multidimensional scaling (NMDS) analysis was run on species communities inside and outside of the halo using data from diver observations at 20 camera locations and species counts from 54 additional images taken in the 15-minute intervals before, during, and after the diver observations at 18 of those locations. Only images from immediately before, after, and during diver observations were used for comparison in order to control for natural variation in fish communities over time, since the main goal of the comparison was to assess the ability of the cameras to capture known community composition at a given location and time. All fish in this subset of images were identified to the genus or species level by the same researchers who conducted the in-water diver observations. To determine whether divers had any effect on fish presence, counts from images captured before, during, and after diver observations were compared using a nonparametric Friedman rank sum test and a Wilcoxon signed rank test for pairwise comparisons. All analyses were conducted using R statistical software version 4.0.3 (R Core Team 2020) with packages qdap (Rinker 2019), reshape (Wickham 2007), rstatix (Kassambara 2020), and vegan (Oksanen et al. 2018).

### Image Analysis

#### Image Sorting

Images were initially named and sorted by location, time, and date using the camtrapR (Niedballa et al. 2016) package in R (R Core Team 2020). A subset of over 13,000 images were sorted by trained undergraduate student volunteers into categories containing at least one visible fish (“Fish”) and without any visible fish (“NoFish”). To ensure consistency, all volunteers were initially trained on the same subset of ~300 images drawn from multiple different cameras and their accuracy assessed by the research team before they were assigned a larger subset of images to sort individually. Due to the nature of timed rather than motion or heat triggered photos, many images did not contain fish.

#### Model Choice

In order to streamline future analyses of fish photos and identification and avoid manual redundancy in discriminating fish existence in pictures, we built and trained a Convolutional Neural Network (CNN) based on ImageNet pretrained ResNet-50 (He et al. 2016), a deep residual network widely used in image classification. We chose ResNet as our base structure because we wanted conservative results in classifying fish pictures. ResNet is powerful at preventing overfitting, making it less likely to omit pictures containing fish. The reproducible code was implemented in Pytorch (He et al. 2016, Paszke et al. 2017, Howard et al. 2018) to identify images with animals (Fig 3).

**Figure 3.**
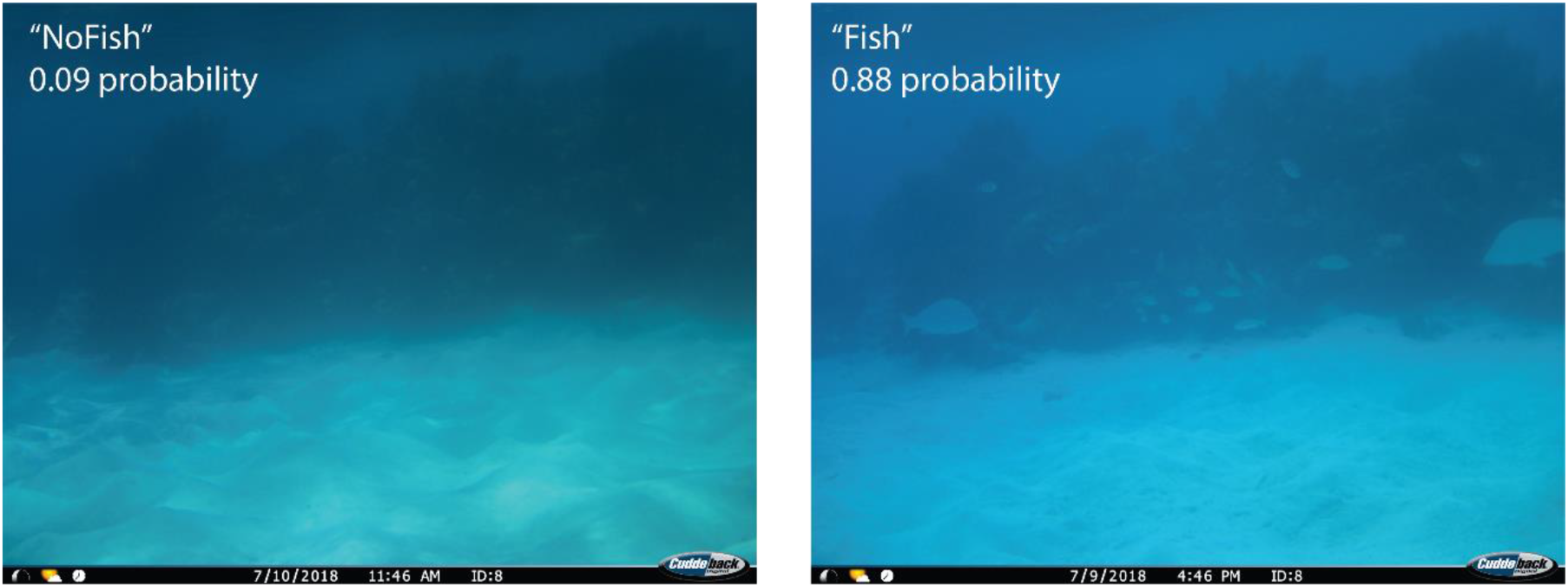
Empty (left) and fish-filled (right) images captured at the same camera site within 24 hours. The empty or “NoFish” photo was assigned only a 0.09 probability of containing a fish by our ResNet-50 model, whereas the model predicted a 0.88 probability of the second photo containing at least one fish, hence its “Fish” designation. Probabilities like these were used to sort images into “Fish” and “NoFish” categories to streamline further analysis.

#### Training and Validation

Our model was trained on a sample of 10,727 images sorted by our team of trained volunteers and then run on 75,470 additional images to sort them into “Fish” and “NoFish” categories. To augment the training set, we preprocessed the fish images with spatial transformations, cropping and lightening variation, and used focal loss (Lin et al. 2017) as an objective function to address the problem of imbalanced labels.

#### Image Analysis

Images with a “Fish” probability of 0.4 or above were used to determine the relative presence of fish at camera sites inside and outside of the Half Moon Caye MPA. A cutoff of 0.4 was chosen based on the model’s reduced ability to correctly identify fish presence, compared to fish absence. This analysis of fish occupancy was conducted based on the relative proportions of the total images classified that did or did not contain fish inside and outside of the MPA and at patch reef and seagrass/algae locations.

## Results

### Camera Performance in the Field

#### Cameras and Housings

Housing 1 proved vulnerable to flooding at depths exceeding 4 m and prone to leaking even at shallower depths. This appears to be due primarily to the thin nature of Housing 1’ s walls and lid, which deformed substantially under pressure, breaking the epoxy seal with the lens and allowing water to enter. Housing 2 proved far more robust to pressure and had minimal leakage even at a maximum deployed depth of 7 m over multiple months. Further testing with this housing will be necessary to determine its maximum functional depth, but preliminary tests with a revised housing design have reached depths of 40 m.

The cameras themselves proved resilient to flooding and were typically still armed and taking photos when extracted from partially-flooded housings. However, many cameras recovered from flooded housings were unable to be redeployed due to lasting damage to their internal systems. Batteries were sufficient to power the cameras past the five month extraction date of our longest deployment, with all non-flooded cameras still displaying the starting battery status of “OK” (not “LOW” or “DEAD”). Maximum water temperature measured at any of our deployment sites, which could affect battery life, was 30° C (86° F). Cuddeback advertises that their cameras can operate up to 12 months continuously with efficient battery usage. Memory cards were more than sufficient to store images on the schedule that they were collected during this period (approximately 8.5 GB of images over 5 months), suggesting that image frequency could be doubled or tripled in future deployments, assuming that processing of these additional images is sufficiently streamlined with the use of computer vision or similar techniques.

#### Environmental Variability

Different benthic environments and camera positioning affected the quality and interpretability or classification of images. In shallow seagrass and macroalgal environments, suspended organic matter in the water column reduced visibility, especially at certain times of day under specific light conditions. Suspended organics may also have contributed to high biofouling in these benthic environments (discussed below). Camera orientation with regard to compass direction was varied between sites during deployment, although both cameras at any given location were oriented in the same direction. While visibility due to the direction and angle of sunlight varied throughout the day at all locations, some orientations produced a greater number of poorly-lit images or images with strong light reflection off of suspended particles in the water column, reducing the overall image quality or number of usable images for those sites.

#### Biofouling

The most significant impediment to long-term camera function at our sites was biofouling, the growth of marine organisms over the camera housing and stands. Image quality declined rapidly due to biofouling, making fish identification impractical in as little as one month depending on site conditions (Fig S2), with a median value of two months endurance, although imagery from some cameras remained usable up to four or five months after deployment. This biofouling decreased both fish detections in images and the ability of our model to accurately classify images taken by biofouled cameras (Fig S3, S4). Algal grazing by fish was an important factor reducing fouling on cameras and stands placed near patch reefs, while cameras placed in seagrass or algae beds away from patch reefs were overgrown with algae over the same deployment period that halo cameras remained relatively unobstructed (Fig 4). The addition of antifouling boat paint to the camera housings before the August 2019 deployment appeared to be only a minor deterrent to organisms growing on the housing in general and did not prevent the camera lens window from being almost completely obscured 2-3 months after deployment. In addition to biofouling, camera function was also limited by reduced visibility at certain sites, which varied based on local turbidity and light availability, and by depth. Preliminary tests of a similar camera design in Hawai’i suggest that the extreme biofouling observed at Lighthouse Reef is not representative of all such reef environments and may be related to especially high productivity and suspended organic matter in the water column at this location.

**Figure 4.**
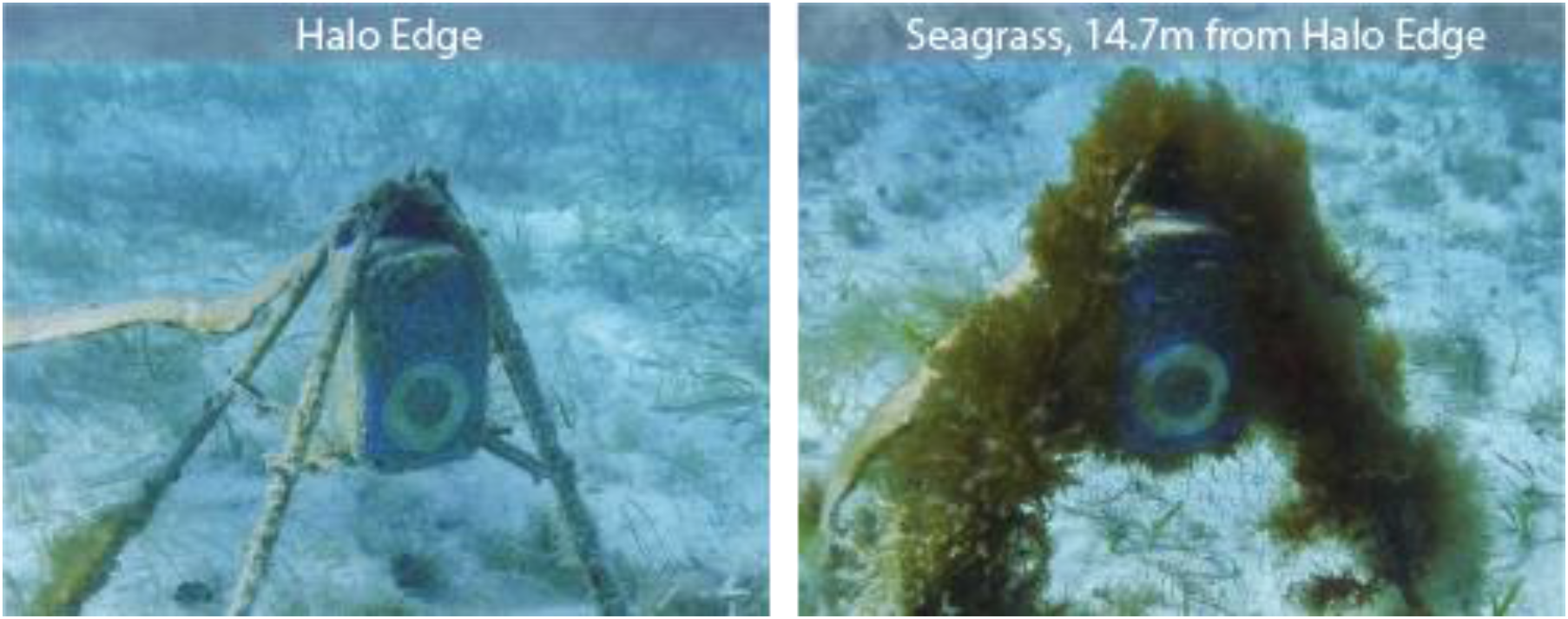
Comparison of biofouling and algal overgrowth of structures placed at the edge of the halo, adjacent to seagrass (left), and in middle of the surrounding seagrass (right) after a month. Similar results seen across all camera sites suggest that reef-based herbivores help to control algal growth both within and at the edges of the halo but do not graze heavily, if at all, on algae and related marine organisms in the surrounding algal or seagrass beds.

### Image Classification and Accuracy

Of the 130,621 images collected, 88,899 were deemed usable (68%) based on a visual examination, despite some level of biofouling in many of these. Our ResNet-50-based model had an overall accuracy of 92.5% when classifying these images into “Fish” or “NoFish” categories, with higher accuracy in identifying empty or “NoFish” photos due to the increased number of these in the training dataset (Table 1).

**Table 1.** Accuracy of model predictions. Accuracy is determined by comparing the computer’s prediction for each image with the actual label of “Fish” (image contains at least one fish or similar animal) or “NoFish” (image is empty) assigned to the photo by a team of trained volunteers. The model was trained on 10,727 images classified by volunteers and then validated with an additional 2,702 human-sorted images, the results of which are shown in this table. Prediction accuracy is higher for “NoFish” (empty) images, likely due to the higher proportion of this type of image in both the training and validation datasets.

| True Image Designation | Correct Predictions | Incorrect Predictions | Percent Accuracy (%) |
| --- | --- | --- | --- |
| *Fish* | 444 | 165 | 72.9 |
| *NoFish* | 2057 | 36 | 98.3 |
| **All images** | **2501** | **201** | **92.5** |

### Community Composition

#### Comparison with Diver Observations

Large compositional differences were identified between the halo/reef and seagrass/algae fish communities but no obvious difference between the communities constructed from diver species observations and those from camera images (NMDS; Fig 5), although individual images consistently contained fewer fish than were observed by divers during a 15-minute period at the same site (Fig S5). An ANOSIM conducted on the NMDS output showed a significant difference between the halo/reef and seagrass/algae communities (R=0.66, p<0.001), which was reflected in both the camera and diver species observations. Overall, there was a small but significant effect of time (before, during, or after diver presence) on fish counts (Friedman rank sum test p=0.0089, Kendall’s W = 0.24). Fish counts were significantly higher during diver observations compared to before (WSRT adj. p=0.021), but there were no significant differences between the before and after time points (WSRT adj. p=0.83) or during and after (WSRT adj. p=0.34), which suggests that divers did not have a consistent measurable effect on fish presence during these short observational periods.

**Figure 5.**
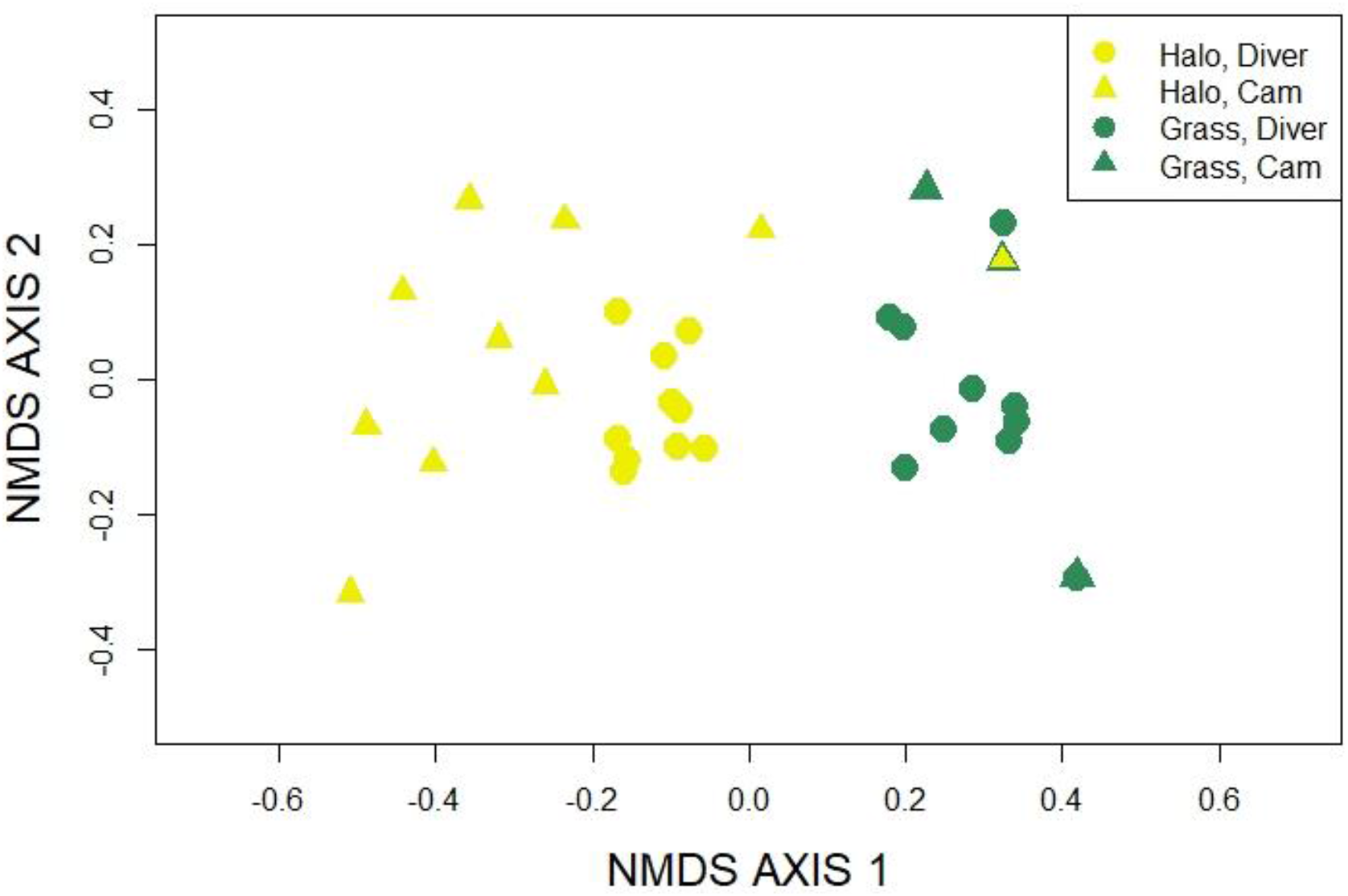
Non-metric multidimensional scaling (NMDS) based on 20 diver observations and an additional 54 camera images from the same sites. Points represent individual site observations or images and are colored yellow for halo/reef cameras and dark green for seagrass or algae cameras. ANOSIM of this NMDS output found a significant difference between halo/reef (yellow) and seagrass/algae (green) communities (R=0.66, p<0.001). Dots represent diver observations, while triangles represent camera images. Most image - based points fall within the same area as those from diver observations, illustrating that the cameras accurately captured the two different fish community types.

#### Fish Distributions

The fish detection counts based on camera data revealed that the proportion of images with fish was 4.8 times higher in halos (0.38 ± 0.002) as compared to seagrass or algae camera locations (0.08 ± 0.002). MPA status also had a significant effect, with an 18% increase in the proportion of photos with fish within the MPA (0.26 ± 0.002), compared to outside the MPA (0.22 ± 0.002). Detailed analysis of camera locations relative to both benthic cover (i.e. reef/halo or algae/seagrass) and protection status (inside or outside MPA) revealed that fish detection is higher in halos outside of the MPA, relative to inside, but lower in seagrass communities outside of the MPA, relative to inside (Fig 6).

**Figure 6.**
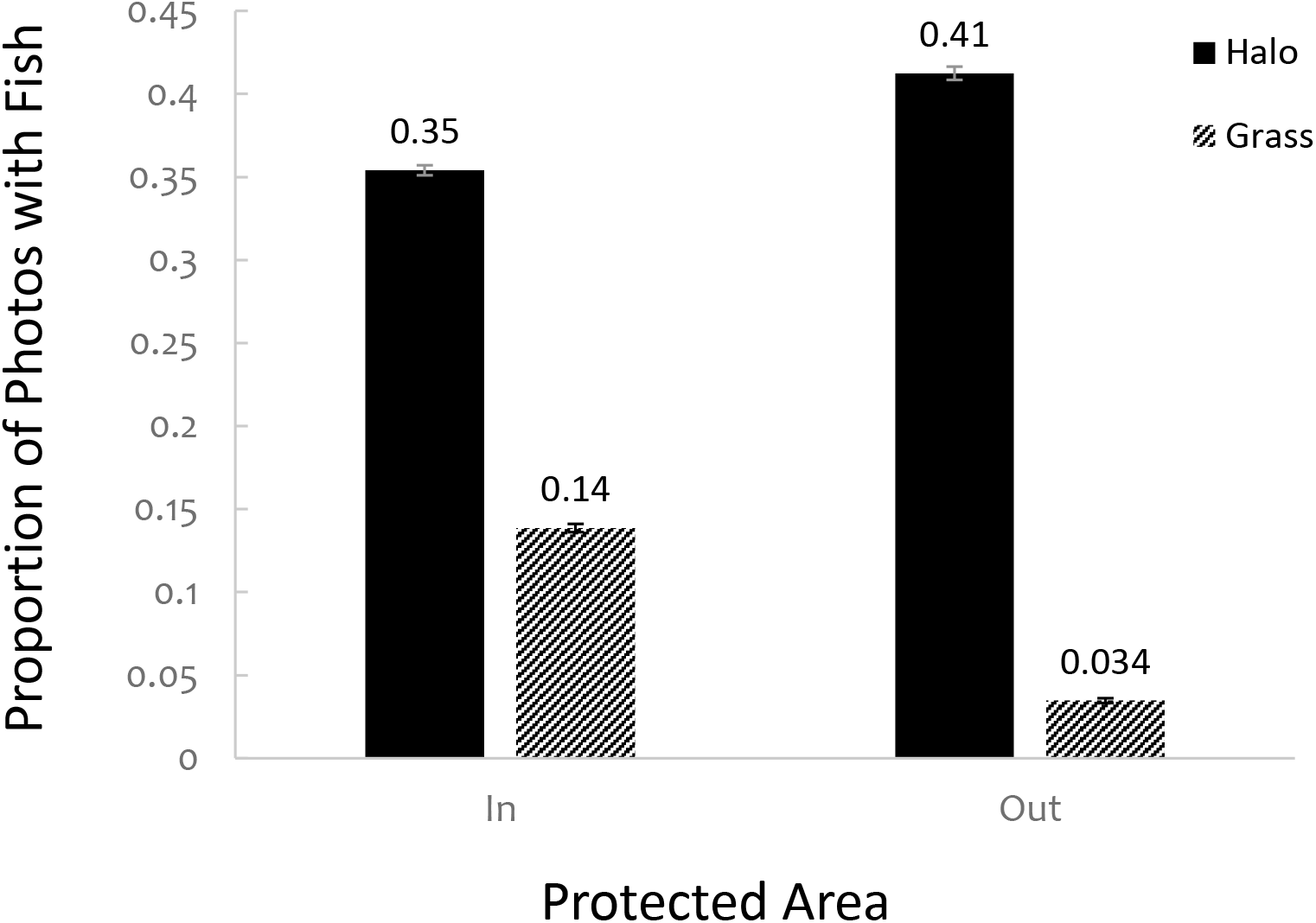
Proportion of images containing fish at halo and grass/algae sites inside and outside of the Half Moon Caye MPA. Proportions calculated to two significant figures are indicated above each bar. Error bars represent ± 1 standard error of the mean from the binomial distribution (≤ 0.004).

A preliminary analysis of diver observations from 20 camera locations as well as an additional 60 images from those sites did not contain a single instance of an herbivorous reef fish (surgeonfish and all adult parrotfish except for *Sparisoma radians*) outside of a halo.

## Discussion

### Camera Performance

#### Deployment Duration and Function

Underwater camera traps proved to be energy-efficient, durable, and capable of producing large volumes of quality images representative of the fish communities at their locations. The DEAC camera trap design is a reliable, cost-effective, and easy-to-implement solution allowing the expansion of terrestrial camera trapping techniques to shallow marine environments. The set-up can easily be used to vastly expand the capability of BRUVs and associated techniques (Cappo, Speare, & De’ath, 2004; Colton & Swearer, 2010; Brooks, Sloman, Sims, & Danylchuk, 2011), and can also provide marine observations over periods of months. In addition to being long-term and more scalable, un-baited camera traps such as these may be more accurate than BRUVs in their estimation of population and community composition (Mccoy et al. 2011) and allow the use of random encounter models for occupancy that are widely used with terrestrial camera traps (Rowcliffe et al. 2014).

Camera placement and orientation is an important consideration when deploying DEACs in the field, and the ideal placement may vary with geographic location, season, target time of day, water quality, and shading from local structures like patch reefs or docks. The cameras have no observable effect on the behavior of marine animals and may therefore be used to document species that typically evade divers. Even cameras that are “cleaned” by herbivorous fish in the halo do not appear to attract any more attention than natural patches of algae or coral rubble. They are also capable of remaining underwater at depth for long periods and recording high volumes of photo or video observations—our longest running cameras were deployed for 5 months, captured over 7,500 images each, and still had battery life to extend to a year. The exceptional duration and the ability to capture vast numbers of images, and to automatically recognize photos with fish (see below) gives the capability to not only provide a continuous record of rare species behaviors or visits by transient species that divers would miss or only observe by chance, but to also, for the first time, to record changes in behavior, relative abundance, and migrations across seasons and lunar phases.

#### Comparison to Existing Methods

DEACs offer significant advantages over existing underwater camera options, as well as surveys by divers. These underwater camera traps record similar data to that gathered by in-person observations, with a few notable exceptions. While the longer time interval chosen for our extended duration field tests did not capture as many individual fish in each image as divers recorded during the same 15-minute period corresponding to a single photo, the cameras did accurately capture the composition of the community during that time, with regard to both species presence and abundance (Fig 5). Analysis of synchronous diver and camera observations at different sites reflected obvious differences in community composition between different benthic environments. These compositional differences also support the placement of the halo and control cameras at each site, suggesting that the cameras were separated enough to capture these different adjacent community types while remaining close enough to control for local site conditions. The only group of animals observed by divers that were not well captured by DEAC images were schools of roving juvenile fish and small parrotfish that camouflage well within seagrass and algal environments and are best detected through movement. The inability of the camera images and corresponding CNN to identify these fish may be due to a combination of more limited image quality in these turbid environments and the inability of human sorters (on whose data the CNN was trained) to distinguish these small fish later in a static image. Since these species are neither significant grazers (in terms of biomass consumed) nor threatening predators, their detection or inclusion in fish community composition was relatively unimportant for addressing the question used here to assess DEAC utility. However, for applications where detection of these species or similarly small or cryptic organisms is important, use of short video clips in place of still images (an option readily enabled with the DEAC design) could increase the ability of both humans and machines to detect these animals through motion.

The current duration of camera operation is over an order of magnitude longer than the operational duration achieved by other non-tethered underwater cameras (e.g. Williams et al., 2014; Siddiqui et al., 2018), even with biofouling seriously impairing image quality after 1-2 months (Fig S2). To the best of our knowledge, these are the first underwater camera traps to be affordable, power-efficient (therefore deployable over the span of months) and also self-contained, without the challenges imposed by a surface-tethered external power source, which typically limits deployment in remote regions and at multiple sites over large areas.

#### Future Improvements

The greatest immediate limitation to the camera design was the inability to prevent or seriously reduce biofouling without periodic manual cleaning of the cameras, although the magnitude of this challenge may be location-specific. Biofouling is a common problem with unsupervised underwater monitoring equipment (Delauney & Compère, 2009), with multiple solutions proposed to control it, including local chlorination (Delauney & Compère, 2009; Xue et al., 2015), copper sheeting and mesh, and UV radiation (Patil, Kimoto, Kimoto, & Saino, 2007). Ongoing tests of camera and housing design are incorporating these methods to reduce biofouling by marine organisms and extend the functional life of the lens window to better reflect the power and storage capabilities of the camera.

While both the batteries and memory cards we employed were sufficient for the deployment duration and image capture frequency tested, the inability to utilize memory cards with greater than 32GB of storage in Cuddeback cameras is problematic with a more frequent time lapse interval or in the case of short videos collected in place of single images. Therefore, optimizing and expanding both power and data storage capacity of these cameras via commercial means (cameras with larger SD storage capability) or noncommercial modifications is another potentially valuable direction for future research. For example, SD Ultra Capacity (SDUC) storage media currently affords 2 TB of storage, vastly expanding the capability of underwater camera traps for either smaller time intervals between images or multi-year deployments. Indeed, large storage capacity may obviate the need for camera triggering, making time-lapse or video with on-board object recognition a superior alternative, giving the ability to capture even rare or transient species while effectively using the millions of potential images generated.

Design improvements currently in progress focus on reducing biofouling, extending the depth range of the camera housings to over 50 m, and implementing an external flash or other nighttime illumination.

### Image Analysis

The success of our initial image sorting using the CNN ResNet-50 illustrates the power of machine learning and computer vision techniques to drastically reduce time and cost when dealing with large image sets. Our ability to use this trained model to pre-sort images with high accuracy before attempting further analysis via manual or autonomous machine-learning based methods also reduces the cost of the most disadvantageous aspect of our timed image capture method: the number of frames with no objects of interest to the current study. Now that datasets obtained from this and similar shallow marine environments can be easily sorted to exclude non-target images, future underwater camera trap projects using a similar time-lapse method will be able to quickly remove the majority of empty frames, while retaining the ability to measure frequency of detection events and variations in fish presence by time of day, season, or other environmental variables by comparison of occupied and empty images. Expansion of the deep neural network model to focus on identification of individual species, functional groups, and/or the sizes of different individuals, as has been done in similar image analyses (Boom et al. 2014, Boussarie et al. 2016, Siddiqui et al. 2018, Norouzzadeh et al. 2018), will further streamline the analysis of this and related large image datasets.

Our community composition analysis also demonstrates that the 15 minute photo interval of our cameras was sufficient to capture species representative of the same fish community observed by divers in the water. This suggests that while a single image does not capture every fish active in the area during the 15 minute period it represents, the collection of images from any given site are representative of the community at that location and are likely to accurately reflect changes in species behavior, abundance, or diversity at the site over a range of time scales (e.g. daily vs. seasonal changes). Further study of camera captures vs. diver observations regarding species known to be wary of divers, either those using traditional open-circuit SCUBA or closed-circuit rebreathers, is an important line of future investigation to understand true reef fish occupancy and abundance.

Our analysis of the number of images with fish collected at different groups of cameras revealed a difference undetected by the fish counts from our diver observations, showing that fish were detected by cameras more frequently (and are therefore likely to have higher occupancy) in halos outside of the MPA, while fish were detected in seagrass or algal habitats more frequently inside of the MPA. The results of this apparent interaction between benthic cover and protected status illustrate the complexity of patterns and variation within the benthos and the fish communities at Lighthouse Reef.

Further image analysis and modeling may help to distinguish the underlying causes of such variation. The inability of traditional diver surveys alone to detect these differences at all reinforces the value of spatially-distributed, long-term datasets like the images collected from our cameras. The observed differences in fish detection between halo and grass/algae control sites also reinforce diver observations of both less fish and a different fish community in the algae or seagrass beds away from the reef and support the results of the community composition analysis (Fig 5).

### Case-Study and Future Applications

We obtained a large volume of usable images from sites with a variety of depths and benthic cover types, subjected to different fishing pressures inside and outside of a local MPA, and monitored over the course different seasons. This allows us to reasonably conclude that our observations are likely representative of fish presence and behavior at patch reef sites within Lighthouse Reef Atoll as a whole. The complete lack of grazing reef fish observed outside of the halo region by cameras located in surrounding sea grass within 30 m or less of the halo supports the idea that a predator modification of prey fish behavior imposes real constraints on fish movement and that heightened grazing pressure adjacent to reef structures is not simply the result of fish randomly dispersing with distance from the reef. If the latter were true, the pattern would be a simple exponential decrease in fish as distance from the patch reef increased, which would likely be reflected in a lower (but nonzero) number of grazing fish detected at seagrass and algae camera locations, compared to those at the halo edge. An alternative hypothesis is that the lack of grazing fish appearing in images taken outside of the halo is due to unequal detection of fish between the two environments, possibly the result of faster or more furtive movements outside the relative safety of the halo. However, this explanation is refuted by two pieces of evidence: First, diver observation also showed no herbivorous reef fish at grass or algal camera sites, and, second, other fish species that were observed by divers to move quickly through the seagrass or algal environment (e.g. bar jacks, *Caranx ruber*) appear in both halo and control camera images, indicating that cameras placed in the grass are very capable of photographing fish moving through that environment. It is therefore more likely that the complete lack of detection of reef-based grazers outside of the halo, even though their food is in much higher supply there, is due to their total or near-total absence from this environment because of the combined lack of shelter and exposure to predators.

This study demonstrates the value of long-term, spatially-distributed underwater camera trap observations for addressing a subtle difference in fish communities and benthic pattern generation. DEACs can be easily deployed alone or in large spatial arrays for short or extended time periods, and they are capable of recording periodic still images for long-term studies or short videos for detailed behavioral observations. Most importantly, these cameras are highly energy-efficient and require little-to-no maintenance while deployed, making them ideal for remote locations or extended observations that surface-anchored systems or commercial underwater cameras with limited battery life are not suitable for. Multiple networks of camera traps like TEAM (Beaudrot et al. 2016) and Snapshot Serengheti (Norouzzadeh et al. 2018) have been successfully deployed over large regions in terrestrial environments, and DEACs offer the option to now expand such long-term, spatially-extensive monitoring efforts to the marine realm. The use of a CNN makes processing the volume of images collected over such a long-term study a practical option, and continuing advancements in machine learning and computer vision are likely to enable further processing of similar large visual datasets in the future.

## Supporting information

All Supplemental Figures

## Acknowledgements

We thank the Belize Audubon Society for hosting us at Half Moon Caye and assisting with data collection, especially Eli Romero, Dominique Lizama, and Shane Young. This work was conducted under Belize Fisheries Permit 00021-18 and was partially funded by the Wake Forest Center for Energy, Environment, and Sustainability, a Wake Forest University Richter Scholarship, Sullivan Scholarship, and Undergraduate Research Fellowship to Austin Schwartz and a Wake Forest University Vecellio Grant to Stephanie Bilodeau. We thank Greg Asner and the Carnegie Airborne Observatory/ASU Center for Global Discovery and Conservation Science for use of their depth models. We also thank Elvis Solis for logistics help and counsel, Connor Walsh and John Gorelick for help in the field and in the lab, and all undergraduates who contributed to classifying and analyzing the data, especially Tatianna Stroud and Joseph Chen.

## Author Contributions

SB and MS conceived the ideas and designed the camera trap methodology. BX and VPP designed and trained the machine learning model. SB, AS, and MS collected the data. SB and BX analyzed the data. All authors contributed to the drafts and gave final approval for publication.

