## Supplementary material for "A low-cost, long-term underwater camera trap network coupled with deep residual learning image analysis": All Supplemental Figures

### SUPPLEMENTAL DATA

AUTHORS: Stephanie M. Bilodeau[1,2], Austin W. H. Schwartz[1,2], Binfeng Xu[3], V. Paul Pauca[3], and Miles R. Silman[1,2]

INSTITUTION:

1. Department of Biology, Wake Forest University, 1843 Wake Forest Rd, Winston-Salem, NC 27109
2. Center for Energy, Environment, and Sustainability, Wake Forest University, 1843 Wake Forest Rd, Winston-Salem, NC 27109
3. Department of Computer Science, Wake Forest University, 1843 Wake Forest Rd, Winston-Salem, NC 27109

### SUPPLEMENTARY FIGURES

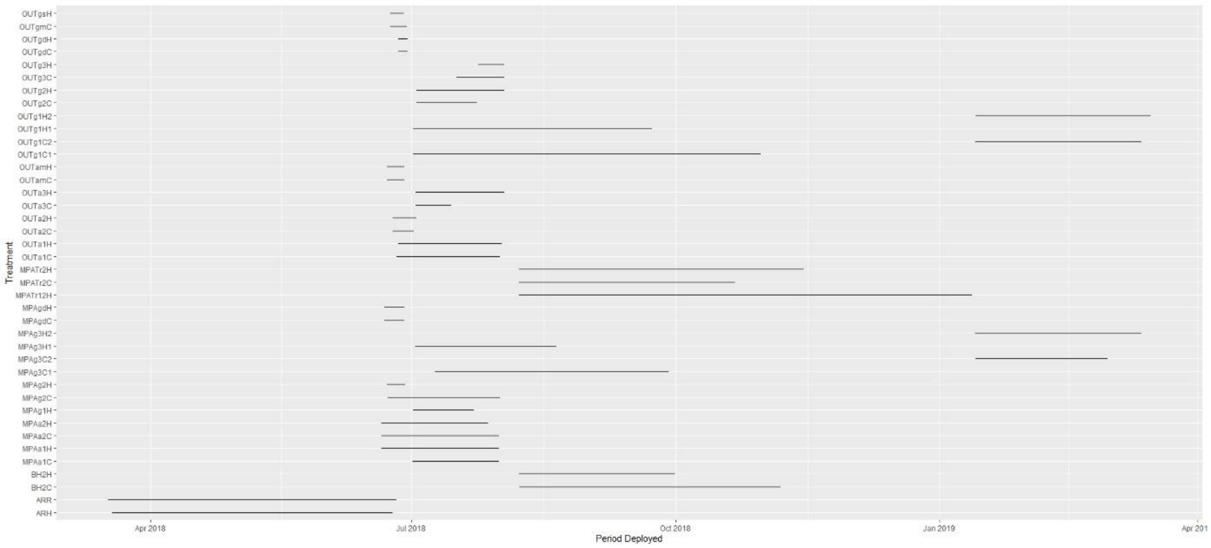


***Figure S1.*** *Deployment periods for individual cameras, typically deployed in pairs at a given site. Treatment designations represent if the camera was deployed in the Half Moon Caye marine protected area (MPA) or outside (OUT), in algae (a) or seagrass (g), the site number or other site designation within that treatment, and camera location at the halo edge (H) or farther away (C),. The final number present for only certain treatments indicates which of multiple deployments at the same location this period refers to.*

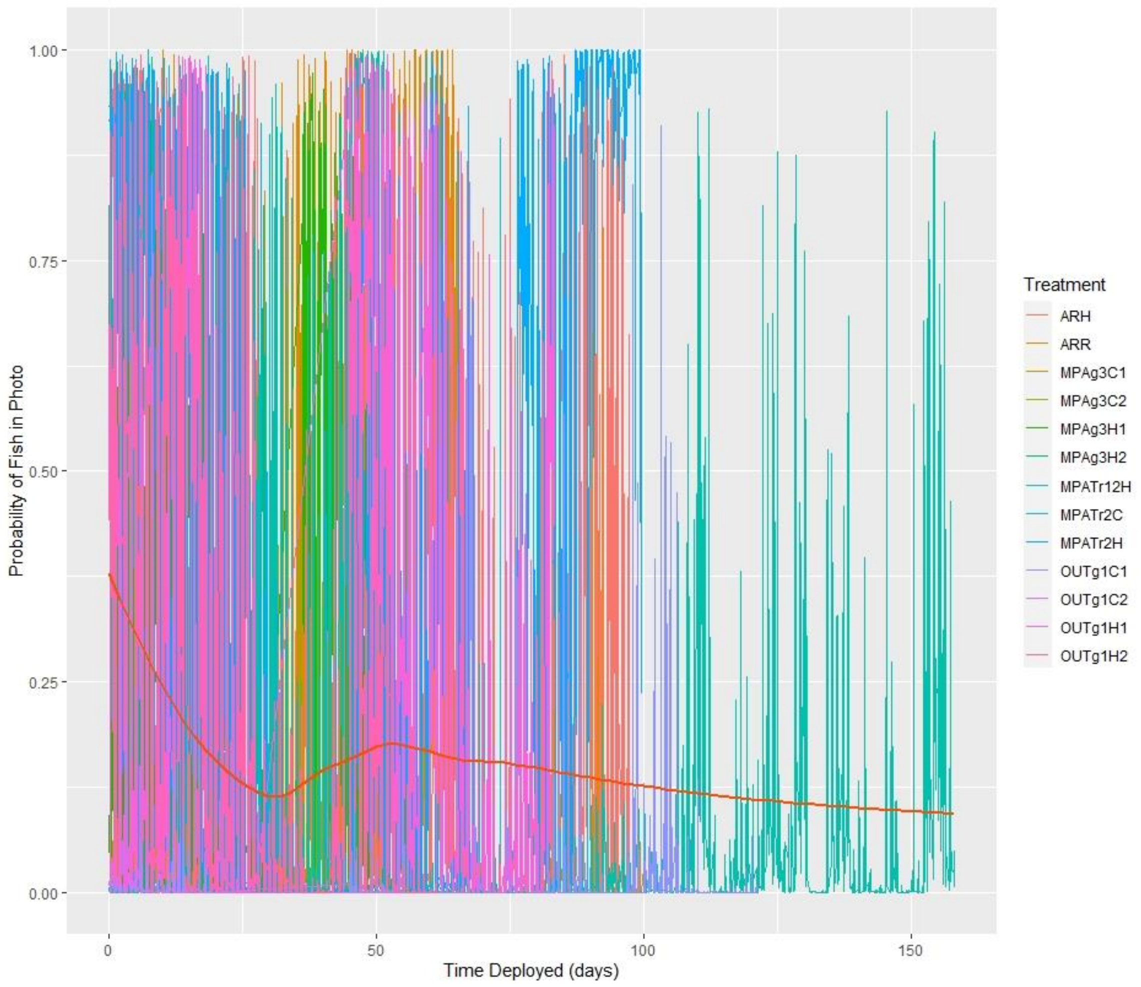


***Figure S2.*** *Decline in fish detections due to biofouling over time for the 13 longest deployments. Deployment time in days is shown on the x-axis, while the probability of a given image containing at least one fish is shown on the y-axis, as calculated by the ResNet-50 model. Fish detection probabilities decrease over the first month, at which point the pattern becomes less clear, likely the result of inaccurate image classifications at certain sites in response to biofouling in the images. The red line represents the smoothed results of all sites combined, using the "loess" function in R package ggplot2 (Wickham 2009).*

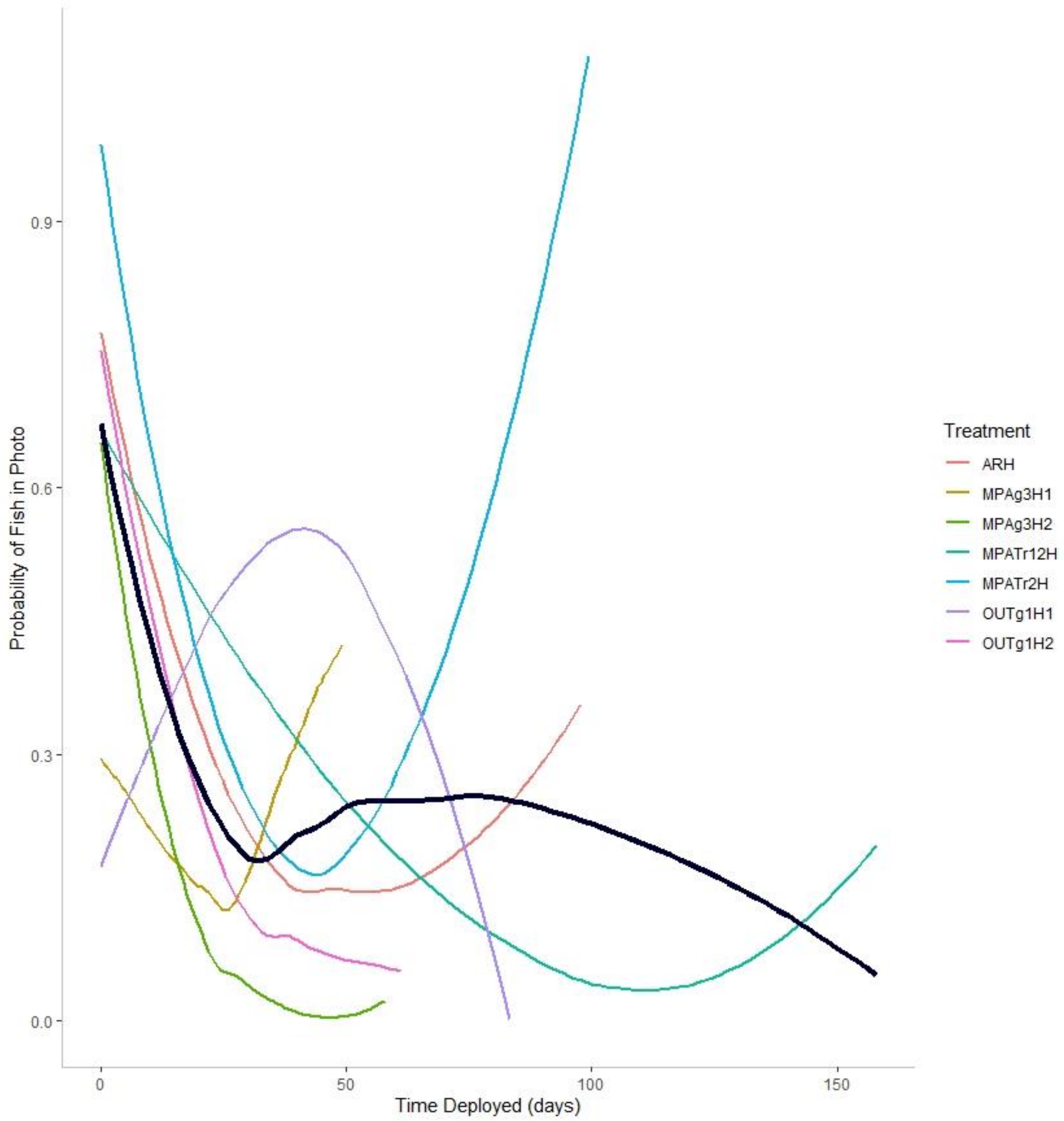


***Figure S3.*** *Fitted lines for fish detection probabilities over long-term reef-adjacent halo deployments. Lines were fitted with the "loess" local polynomial regression fitting function with span = 1 from R package ggplot2 (Wickham 2009). The dark line represents the smoothed results of all sites shown on this graph. Not only do fish detections decline consistently at most sites due to biofouling of the camera lens over the first month, model accuracy is clearly impaired past this point at certain sites, leading to a false increase in fish detection probability for some cameras (e.g., note the erroneously high probabilities around 80-100 days for MPATr12H, shown in blue).*

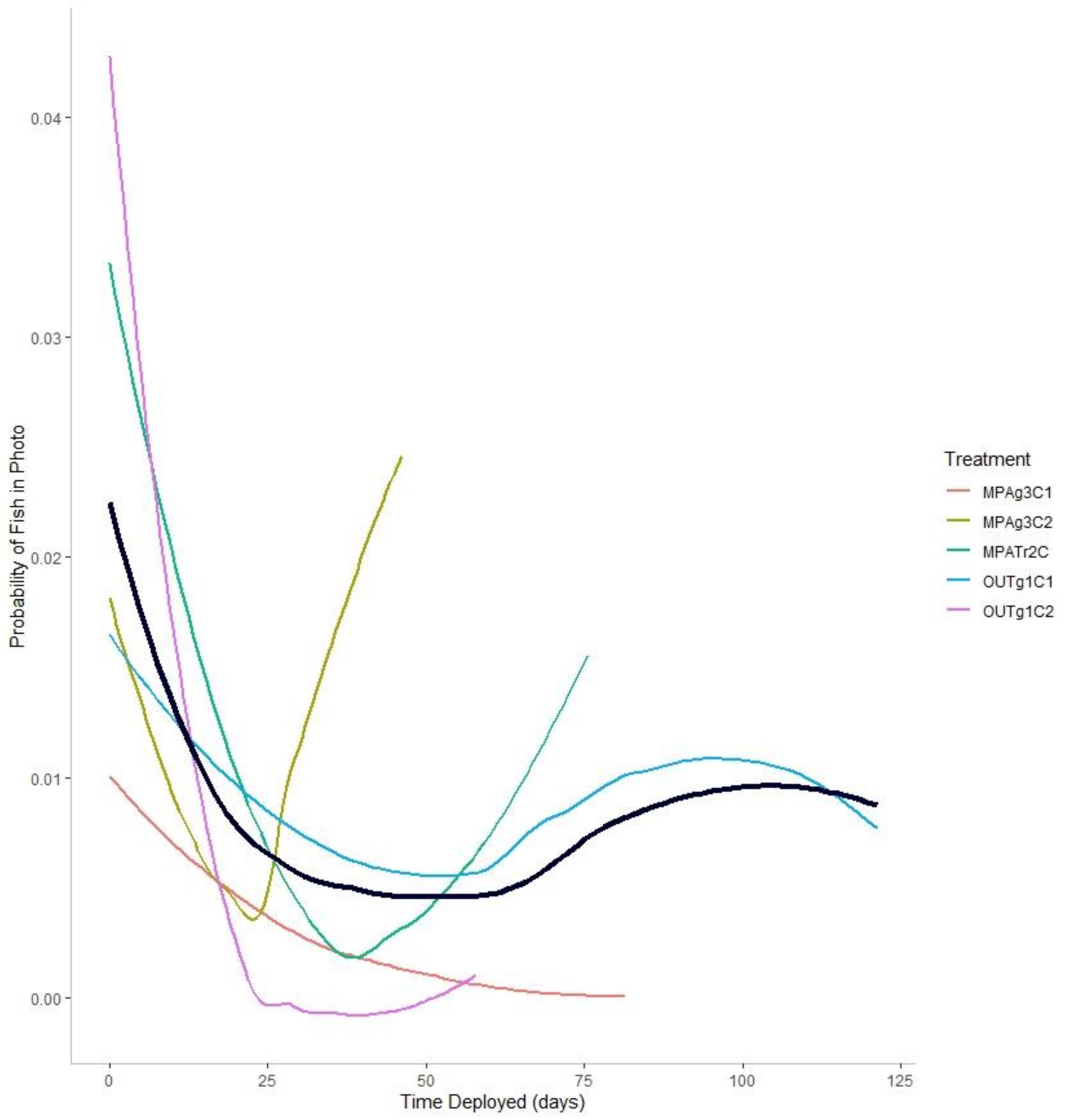


***Figure S4.*** *Fitted lines for fish detection probabilities over long-term seagrass or algae meadow deployments. Lines were fitted with the "loess" local polynomial regression fitting function with span = 1 from R package ggplot2 (Wickham 2009). The dark line represents the smoothed results of all sites shown on this graph. Note that fish detection probabilities are lower at grass sites, relative to reef/halo sites, and the y-axis is scaled appropriately. After the first month, fish detection probabilities erroneously rise at a couple sites, likely reflecting reduced accuracy in the ResNet-50 model's image classifications as a result of biofouling.*

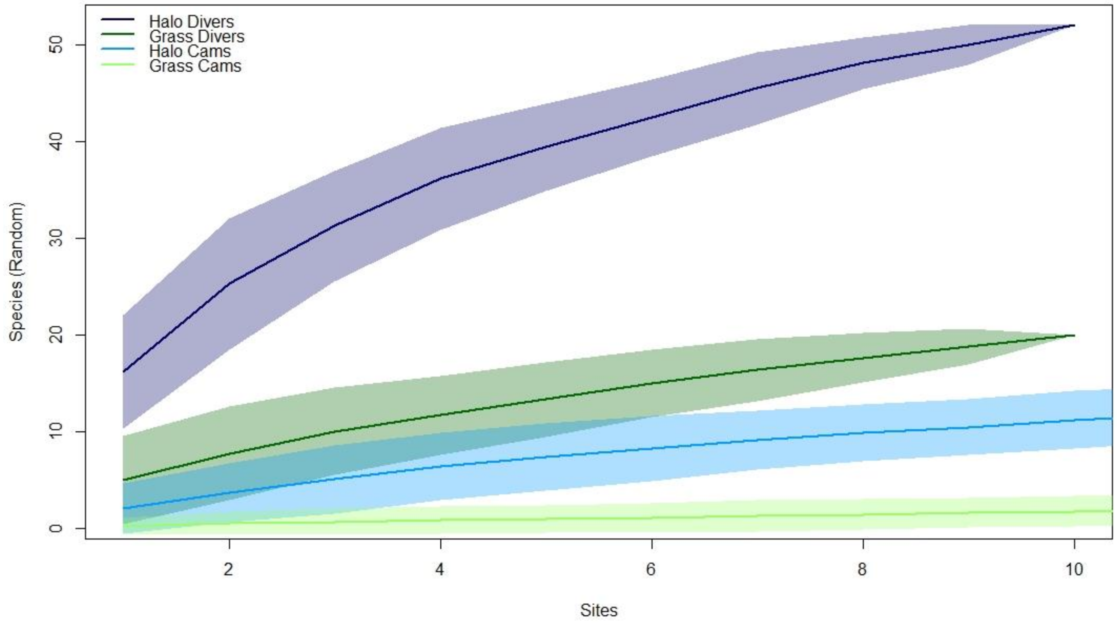


***Figure S5.*** *Species accumulation curves from 15-minute continuous diver observations and camera images from the 45-minute window surrounding each observation (3 images for each site). Divers clearly observe more species than cameras over these short periods, but the overall community compositions constructed from these species counts are similar between divers and cameras (see Fig 5).*
